# Contrasting patterns of divergence at the regulatory and sequence level in European *Daphnia galeata* natural populations

**DOI:** 10.1101/374991

**Authors:** Suda Parimala Ravindran, Maike Herrmann, Mathilde Cordellier

**Affiliations:** Universität Hamburg, Institute of Zoology, Martin-Luther-King Platz 3, 20146 Hamburg, Germany; Department of Veterinary Medicine, Paul-Ehrlich-Institut, Paul-Ehrlich-Straße 51-59, 63225 Langen, Germany

**Keywords:** constitutive gene expression, RNA-seq, molecular phenotype, population transcriptomics, DRIFTSEL

## Abstract

Understanding the genetic basis of local adaptation has long been a focus of evolutionary biology. Recently there has been increased interest in deciphering the evolutionary role of *Daphnia*’s plasticity and the molecular mechanisms of local adaptation. Using transcriptome data, we assessed the differences in gene expression profiles and sequences in four European *Daphnia galeata* populations. In total, ~33% of 32,903 transcripts were differentially expressed between populations. Among 10,280 differentially expressed transcripts, 5,209 transcripts deviated from neutral expectations and their population-specific expression pattern is likely the result of local adaptation processes. Furthermore, a SNP analysis allowed inferring population structure and distribution of genetic variation. The population divergence at the sequence-level was comparatively higher than the gene expression level by several orders of magnitude and consistent with strong founder effects and lack of gene flow between populations. Using sequence information, the candidate transcripts were annotated using a comparative genomics approach. Thus, we identified candidate transcriptomic regions for local adaptation in a key species of aquatic ecosystems in the absence of any laboratory induced stressor.

## INTRODUCTION

Natural genetic variation shapes divergence in phenotypic traits and is an important resource for evolutionary processes (Oleksiak *et al.* 2002). Populations respond to environmental variation by genetically adapting to their environments (Hereford 2009; Kawecki & Ebert 2004; Savolainen *et al.* 2013), often showing variations at both gene expression and sequence level across the geographic range of a species. One of the fundamental goals of research in the field of molecular evolution is to resolve the evolutionary processes driving the rise and maintenance of expression and sequence polymorphisms behind this variation. Revealing their effect on an organism’s fitness thereby aids to understand the genetic basis of local adaptation (MacManes & Eisen 2014). Gene expression patterns link genotypes and phenotypes, sometimes called a “molecular phenotype”, and as such is an important component in local adaptation processes (Lopez-Maury *et al.* 2008). --Several studies have reported the testing of different populations exposed to different treatments and examining their transcriptional response, for example in springtails (*Folsomia* (De Boer *et al.* 2013) and *Orchesella* (Roelofs *et al.* 2009)), oyster (*Crassostrea virginica*; Chapman *et al.* 2011; Chapman *et al.* 2009), sparrows (*Zonotrichia capensis*; Cheviron *et al.* 2008), flounder (*Platichthys flesus*; Larsen *et al.* 2008), and seagrass (*Zostera marina*; Jueterbock *et al.* 2016; Reusch *et al.* 2008), thereby identifying candidate genes involved in local adaptation. Gene expression variation can be highly heritable (Brem & Kruglyak 2005; Schadt *et al.* 2003; Whitehead & Crawford 2006b). Moreover, constitutive gene expression patterns also differ within-and among-natural populations (e.g., Roberge *et al.* 2007; Whitehead & Crawford 2006a), strongly suggesting that standing variation in constitutive gene expression is shaped by local adaptation. Natural selection acts immediately on newly arisen variation (in contrast to adaptation observed from standing genetic variation) as there are neutral and slightly deleterious variations preserved in a population, which may become beneficial upon changes in selection regimes (Barrett & Schluter 2008). After a sudden change of environment, standing variation can contribute to fast adaptation (Feulner *et al.* 2013; Kitano *et al.* 2008). Identifying allelic/genetic variants underlying differences in expression profiles can be helpful in hypothesizing gene functions (Jansen & Nap 2001; Kesari *et al.* 2012; Rockman 2008). Although prior knowledge of the specific loci is not a prerequisite to learn about adaptive processes in most cases, identification of genetic features underlying local adaptation is critical in answering fundamental questions about natural selection (Rausher & Delph 2015).

Geneticvariation within and among populations is strongly influenced by their colonization history, and the demographic changes following the primary establishment of a population. Population sizes may vary after colonization across the species based on environmental factors and further colonization (Böndel *et al.* 2015). Colonization events depend on dispersal ability, and dispersal rates strongly differ from gene flow estimates in several species (De Meester *et al.* 2002). This is particularly evident in freshwater zooplankton species, where several studies suggest a high potential for dispersal when populations rapidly colonize new habitats and spread invasively (Havel *et al.* 2000; Louette & De Meester 2004; Mergeay *et al.* 2008). However, genetic studies show that the observed rate of gene flow is much lower than would be expected in organisms with high dispersal potential (Boileau *et al.* 1992; De Meester *et al.* 2002; Thielsch *et al.* 2009). This ambiguity between dispersal potential and rate of gene flow can be explained by founder effects (Boileau *et al.* 1992) complemented by local adaptation; resulting in monopolization of resources by local populations (De Meester *et al.* 2002). This process leads to the impression that population genetic variation correlates with the colonization patterns (Orsini *et al.* 2013).

Amongst freshwater zooplankton species, the water flea *Daphnia* is the best studied and has been an important model for ecology, population genetics, evolutionary biology, and toxicology (Ebert 2005). This genus belongs to the order Cladocera and has attracted scientific interest since the 17^th^ century (Desmarais 1997). It inhabits most types of freshwater habitats and includes more than 100 known species of freshwater plankton organisms (Ebert 2005). *Daphnia* make an interesting subject of investigation in comparative functional genomics (Eads *et al.* 2008). Apart from the fact that *Daphnia* species have an appropriate size for being used in laboratory cultures, they are easy to cultivate and have short generation times. Because of their clonal mode of reproduction, *Daphnia* are highly suited for quantitative genetic studies, which can enhance our understanding of their evolutionary ecology.

Genetic variation has been reported for numerous traits in *Daphnia,* such as life history traits (e.g., Henning-Lucass *et al.* 2016), vertical migration (e.g., Haupt *et al.* 2009), fish escape behavior (e.g., Pietrzak *et al.* 2015), resistance against parasites (e.g., Routtu & Ebert 2015) and immune responses (e.g., Garbutt *et al.* 2014). Furthermore, it was shown that responses to many chemical stressors such as phosphorus (Roy Chowdhury *et al.* 2015; Roy Chowdhury *et al.* 2014), copper (Poynton *et al.* 2008), cadmium (Soetaert *et al.* 2007) and pharmaceutical products like ibuprofen (Hayashi *et al.* 2008; Heckmann *et al.* 2007) have a genetic basis as well. Within-and between-population comparisons in *Daphnia* have been conducted extensively using varied environmental perturbations and providing evidences for local adaptation (for e.g., Barata *et al.* 2002; Declerck *et al.* 2001; Ebert *et al.* 1998; Spitze 1993). Although various aspects like phylogeography, functional morphology, physiology and life history evolution have been in the limelight of *Daphnia* research for several decades (Eads *et al.* 2008), *Daphnia* genomics investigations have begun only in the last decade with the availability of the *Daphnia pulex* genome (Colbourne *et al.* 2011). A considerable number of studies (for e.g.: Bento *et al.* 2017; Miner *et al.* 2012; Orsini *et al.* 2016; Yampolsky *et al.* 2014) on biotic and abiotic factors have been carried out showing how *Daphnia* respond to environmental perturbations by changes in gene expression. However, little is known about the intra-specific variability at the gene expression level in *Daphnia*, since the above-mentioned studies focused on stressor driven responses using a reduced number of genotypes.

To sum up, elucidating the mechanisms by which natural selection acts on gene expression evolutionremains a challenge (e.g.: Fraser 2011; Romero *et al.* 2012). Unraveling the relative consequences of drift versus natural selection on gene expression profiles plays an important role in understanding species divergence and local adaptation. The studies listed above provided evidence for gene expression variation correlated with many environmental factors in *Daphnia*. However, knowledge about the variation in constitutive gene expression structure within and among population is lacking.

In the present study on *Daphnia galeata*, sampled from four different lakes in Europe, we conducted a large scale RNA-seq study in the absence of any laboratory induced environmental stressor. Using transcriptome data, we quantified the constitutive expression profiles and performed a sequence analysis of the four populations. We addressed the following questions: (i) Are there differences in gene expression profiles between the four populations? (ii) How is the observed variation explained by the different levels of organization, i.e., genotype and population? (iii) Do the observed differences in expression profiles result from genetic drift or selection? (iv) What is the role of genetic drift and/or natural selection in shaping sequence variation? (v) What are the functional roles of the transcripts?

Our study brought contrasting patterns of divergence at the regulatory and sequence level into light. While no population specific gene expression patterns were found for majority of the analyzed transcripts, divergence patterns at the sequence level hinted at strong influences of founder effects, bottleneck events and divergent selection. Further, our gene co-expression network analysis showed conserved patterns while assessing the population–specific networks and supported our observations at the regulatory level. We were able to identify candidate transcripts for local adaptation using combined approaches. Further comparative genomics analyses are needed to complement our preliminary functional annotations of these candidate transcripts to identify the ecological drivers behind the observed patterns of adaptation.

## METHODS

### Sampling and RNA collection

A set of *D. galeata* resting stages (ephippia) was collected from the sediment of four lakes: Jordán Reservoir (hereafter, Pop.J) in Czech Republic, Müggelsee in Germany (hereafter, Pop.M), Lake Constance (hereafter, Pop.LC) at the border between Germany, Switzerland and Austria, and Greifensee (hereafter, Pop.G) in Switzerland. These ephippia were hatched under laboratory conditions (see Henning-Lucass *et al.* 2016 for hatching conditions) and the hatchlings were used to establish clonal lines in a laboratory setting. The species identity was checked by sequencing a fragment of the *12S* mitochondrial locus and 10 microsatellite markers (Multiplex 2 comprising the loci *Dgm109, Dp196, Dp281, Dp512, SwiD1, SwiD10, SwiD12, SwiD14, SwiD15, SwiD2*), following protocols by Taylor *et al.* (1996) and Yin *et al.* (2010) respectively.

Mature females for six clonal lines per lake were placed at equal densities (40 individuals L^−1^) in semi-artificial medium for a week, during which the juveniles were regularly removed. Gravid females from the equal density beakers were then collected within three days during a time window of a few hours. Twenty to thirty individuals were homogenized in a 1.5 mL centrifuge tube in 1 mL Trizol (Invitrogen, Waltham, MA USA) immediately after removing the water. The Trizol homogenates were kept at −80 °C until further processing.

### RNA preparation and sequencing

Total RNA was extracted following a modified phenol/chloroform protocol and further processed using the RNeasy kit (Qiagen, Hilden, Germany). The total RNA was eluted in RNAse free water and the concentration and quality (RNA integrity number and phenol) were checked using a NanoDrop spectrophotometer (Thermo Scientific, Wilmington, DE, USA) and a Bioanalyzer 2100 (Agilent Technologies, Santa Clara, CA, USA). The 72 total RNA samples were sent to the company GATC (Konstanz, Germany) for library preparation and sequencing. Following reverse transcription and cDNA construction using random primers, 50bp single-end (SE) reads were sequenced on an Illumina HiSeq 2000 (San Diego, CA, USA), with. To avoid block effects and confounding effects in the downstream analysis, we used a completely randomized design; each library was sequenced on at least two different lanes, on a total of nine lanes. Detailed information can be found in Table S1.

### Quality trimming, mapping and read counts

All reads with ambiguous bases (Ns) were removed before trimming. Bases with a phred score below 20 were trimmed at the 3’ and 5’ ends. Reads shorter than 45 bp after trimming were discarded. All trimming steps were conducted using locally installed version of Galaxy at the Gene Center in Munich, Germany.

Trimmed reads were mapped to the reference *D. galeata* transcriptome (Huylmans *et al.* 2016; available from NCBI: https://www.ncbi.nlm.nih.gov, GenBank ID: HAFN00000000.1) using NextGenMap (Sedlazeck *et al.* 2013) with increased sensitivity (-i 0.8 –kmer-skip 0 −s 0.0). Read counts were obtained from the SAM files using a custom python script (available upon request) and discarding ambiguously mapped reads. The raw count table was analyzed in R (R Development Core Team 2008) using the package DESeq2 (Love *et al.* 2014). Normalization was done with size factor procedure. Standard differential analysis steps of DESeq2 such as estimation of dispersion and negative binomial GLM fitting were applied. The count outliers were automatically detected using Cook’s distance, which is a measure of how much the fitted coefficients would change if an individual sample was removed (Cook 1977). Principal Component Analysis (PCA) was performed to visualize the clustering of biological replicates and clonal lines.

To identify the differentially expressed transcripts (DETs) upregulated the most in each population, we used the DESeq2 “contrasts” function. We performed six pairwise comparisons: Pop.G vs Pop.J, Pop.G vs Pop.LC, Pop.G vs Pop.M, Pop.J vs Pop.LC, Pop.J vs Pop.M, Pop.LC vs Pop.M. All *p*-values were adjusted for multiple testing using the Benjamini-Hochberg correction (Benjamini 1995) implemented in DESeq2. To create a list for each population from each comparison, we retained transcripts that had an adjusted *p*-value (*p_adj_*) equal to or lower than 0.05 and a fold change (FC) deviating from 0 (depending on the direction of the pairwise comparison), resulting in four lists as follows:

1. Pop.G: G vs. M: FC > 0; G vs. LC: FC > 0; J vs. G; FC < 0
2. Pop.J: J vs. G: FC > 0; J vs. LC: FC > 0; J vs. M; FC > 0
3. Pop.LC: J vs. LC: FC < 0; LC vs. M: FC > 0; G vs. LC; FC < 0
4. Pop.M: G vs. M: FC < 0; J vs. M: FC < 0; LC vs. M; FC < 0

The four lists of DETs obtained above were combined to identify population specific transcripts and Venn diagrams depicting the overlap between the contrasts were created using the VennDiagram package (Chen 2011) in R.

### Evaluating the role of natural selection on transcript expression levels: DRIFTSEL

We searched for transcripts for which the identified differential expression could not be explained by phylogenetic distance and genetic drift alone. To identify signals of possible selection, we used the approach of Ovaskainen *et al.* (2011) implemented in the R package DRIFTSEL 2.1.2 (Karhunen *et al.* 2013), considering expression of every single transcript as a trait. To perform this analysis, we made use of the microsatellite data and normalized read count values. Allele frequencies were obtained from microsatellite data collected in a previous study, independently from the species identification step outlined above. Microsatellite data of 30-40 resting eggs also sampled from the same sediment layers the resurrected clonal lines come from was obtained from a study by Herrmann (2017). Briefly, eleven microsatellite loci were analyzed for each individual according to the protocol published by Thielsch *et al.* (2009). Primers for all loci were multiplexed and PCR was performed using the Type-it Microsatellite PCR Kit (Qiagen, Hilden, Germany). Alleles were recorded manually and allelic frequencies were calculated with GenAlEx (Peakall & Smouse 2012).

Using microsatellite allelic frequencies, the coancestry coefficients by admixture F model was calculated using “do.all” function implemented in the RAFM package (Karhunen & Ovaskainen 2012). We ran a total of 200,000 iterations with thinning at an interval of 1,000 and discarded the first 1,000 iterations as ‘burn-in’. The output was a list which contained samples from the posterior distributions of allele frequencies. Values from the posterior coancestry matrix, ‘theta’, were used as input for the Metropolis-Hastings (MH) algorithm along with the normalized read counts for DETs as implemented in DRIFTSEL. We ran a total of 5,000 iterations with thinning at 1,000 samples and discarded the first 100 iterations as burn-in. The output of MH algorithm was a matrix of posterior of subpopulation effects (pop.ef), used to estimate the H.test values. The H.test describes whether the population means correlate with the genetic data more than it would be expected on basis of shared evolutionary history. Large H-values imply that the populations are more locally adapted than expected by chance.

### Intra and inter-population variation

To quantify the respective contributions of the factors “genotype” and “population” to the observed variation in gene expression profiles, we performed an analysis of variance (ANOVA) test in R on the normalized read counts obtained from DESeq2. To correct for multiple testing, *p_ad_*_j_-values were calculated for each transcript using the Benjamini-Hochberg procedure.

### Variant calling and filtering

The variant calling and filtering steps have already been described in Herrmann *et al.* (2017b). Briefly, the aligned reads from RNA-seq data were merged using samtools (Li *et al.* 2009). GATK (McKenna *et al.* 2010) was used to split exon segments, reassign the mapping qualities (SplitNCigarReads) and indels were aligned (RealignerTargetCreator and IndelRealigner). The HaplotypeCaller (DePristo *et al.* 2011) function was used for the initial variant calls for the realigned reads and samples were jointly genotyped using GATK’s GenotypeGVCFs tool. A single vcf file was created and false positive variant calls were filtered with the following criteria: (i) clusterWindowSize = 35; (ii) Quality by depth (QD) < 2.0; (iii) Fisher Strand (FS) > 30.0. This produced a variant dataset with not only biallelic variants but also triallelic variants and indels.

Using the SNPRelate package (Zheng *et al.* 2012) in R/BioConductor, the variant dataset was limited to only biallelic sites for downstream analysis. These were further pruned for linkage disequilibrium considering a threshold of 0.2 (r^2^ > 0.2), thereby retaining 393,514 SNPs. A PCA was plotted using the functions in SNPRelate which include calculating the genetic covariance matrix from genotypes, computing the correlation coefficients between sample loadings and genotypes for each SNP, calculating SNP eigenvectors (loadings) and estimating the sample loadings of a new dataset from specified SNP eigenvectors.

### Neutrality statistics

To obtain alignments of transcript sequences, SNP calling datasets were filtered as described above. Beagle 4.1 (Browning & Browning 2007) was used to phase SNP calling data and a python script (available upon request) was used to parse the phased vcf file to sample sequences in fasta format. After phasing, we obtained 13006 transcripts containing SNPs and the sequences were input in R. A multiple sequence alignment and Tajima’s D statistics (with *p*-values) were obtained population-wise for each transcript using the pegas package (Paradis 2010) in R.

Results from LOSITAN (Antao *et al.* 2008) outlier tests were obtained from Herrmann *et al.* (2017b) to identify loci under selection (see Table S4).

### Heterozygosity and mutation frequencies

The heterozygosity values for the final SNP dataset were calculated with VCFtools (Danecek *et al.* 2011). The ratio between the expected heterozygosity (H_E_) and observed heterozygosity (H_O_) was calculated based on available SNP information and plots were created using ggplot2 (Wickham 2009) in R.

### Sequence vs. regulatory variation

To visualize the proportion of transcripts responsible for local adaptation at regulatory and sequence level, we consolidated the list of transcripts from various analyses as performed above and represented it with an alluvial diagram (http://rawgraphs.io/). In an alluvial diagram, each black rectangle is called a ‘node’, the colored regions linking the nodes are called ‘flows’ and the vertical group of nodes are called ‘steps’. In our analyses, we had four steps: DESeq2, DRIFTSEL, LOSITAN and Tajima’s D.

### Annotation and functional analysis

To functionally annotate the *D. galeata* transcripts, a local sequence alignment using blastn (Altschul *et al.* 1990) against the nr database (downloaded Feb. 2015 via ftp://ftp.ncbi.nlm.nih.gov/blast/db/) was performed. Hits with an eval ≤ 0 and identity ≥ 50% were considered. Additionally, protein domain annotations and orthoMCL (Li *et al.* 2003) results were obtained from (Huylmans *et al.* 2016). Briefly, a search was made for all three *Daphnia* species (*D. pulex, D. magna* and *D. galeata*) using PfamScan (version 1.5) to look into the Pfam A database (version 27.0; Finn *et al.* 2014) together with hmmer3 (version 3.1b; Mistry *et al.* 2013). In order to identify orthologs and be able to compare it to other arthropod species, orthoMCL was used to cluster the amino acid sequences of *D galeata*, *D. pulex* (version JGI060905; Colbourne *et al.* 2011), *D. magna* (version 7; *Daphnia* Genomics Consortium 2015), as well as *Drosophila melanogaster* (version 5.56; St Pierre *et al.* 2014) and *Nasonia vitripennis* (version 1.2; Werren *et al.* 2010) into orthologous groups and determine the inparalogs. Pie charts representing the number of hits obtained for all transcripts and DETs were created using the plotrix package (Lemon 2006) in R.

We classified the orthoMCL clusters into the following categories:

(a) Clusters that contain only *D. galeata*-specific transcripts
(b) Clusters that are shared between *D. galeata* and *D. pulex*
(c) Clusters that are shared between *D. galeata* and *D. magna*
(d) Clusters that are shared between *D. galeata*, *D. pulex* and *D. magna* (*Daphnia*-specific)
(e) Clusters that are shared between *D. galeata* and other arthropods (*D. melanogaster* and *N. vitripennis*)
(f) Clusters that are shared among all five analyzed species (*Daphnia* and both insects)

### Inparalogs and misassemblies

To assess whether *D. galeata* DETs in an orthologous group are “inparalogs”, isoforms or the result of misassembly, we computed the pairwise sequence divergence for those orthoMCL clusters containing DETs from at least two different populations. Since each significantly differentially expressed transcript was assigned as a DET only to the population in which it was upregulated the most, clusters containing more than one DET most likely contained DETs from different populations. Based on the number of populations within their orthoMCL cluster, the DETs were classified into the categories: “1Pop”, “2Pop”, “3Pop” and “4Pop”, and unclustered DETs were categorized as “0Pop”. 0Pop and 1Pop DETs were excluded from further analysis. In total, there were 716 orthoMCL clusters that contained DETs from at least two populations. Pairwise alignments of the amino acid sequences in each orthologous group were performed using the iterative refinement method incorporating local pairwise alignment information (L-INS-i) in MAFFT (Katoh *et al.* 2002). We then used EMBOSS tranalign (Rice *et al.* 2000) to generate alignments of nucleic acid coding regions translated from aligned protein sequences. Pairwise genetic divergence was computed with ‘dist.dna’ function implemented in the ape package (Paradis *et al.* 2004) in R, using the Kimura-2-parameters model with gamma correction. We used an arbitrary cut-off value of 2 to distinguish inparalogs from misassembled sequences.

### Gene Ontology (GO) enrichment analysis

DETs with a H.value ≥ 0.95 (DRIFTSEL result) and transcripts with a nonzero D value in each of the four populations (Tajima’s D result) were analyzed with, “topGO” (Alexa & Rahnenfuhrer 2016) in R, using a custom GO annotation for *D. galeata*. GO terms enriched in the transcripts of interest in each population from each analysis (DRIFTSEL and Tajima’s D) were identified using the ‘weight01‘ algorithm for all three ontologies, namely: molecular function, cellular component and biological processes. We used a Fisher test and those GO terms with a classicFisher value ≤ 0.05 were considered to be enriched for each ontology in each population. A multiple testing procedure was not applied as the *p*-values returned by the ‘weight01’ algorithm are interpreted to be corrected and might exclude “true” annotations (Alexa & Rahnenfuhrer 2016).

### Weighted Gene Co-expression network analysis

To gain insights into the population-specific regulatory patterns of transcripts in *D. galeata*, we performed a weighted gene co-expression network analysis with WGCNA (Langfelder & Horvath 2008) using the variance stabilized normalized read counts obtained in DESeq2 analysis. Transcripts and samples that had lower expression values were excluded from every population using the ‘goodSampleGenes’ function in WGCNA and used for downstream analysis. In total, 32375 transcripts were used for the construction of gene co-expression networks. To identify population-specific co-expression modules (i.e., clusters of highly correlated transcripts), a network was first built using the full dataset (i.e., with samples and transcripts from all populations) and one network for each population using expression values specific to all genotype and biological replicates. The population specific network was compared to the reference network and an adjacency matrix was calculated. Clusters were identified using the WGCNA Topological Overlap Matrices (TOM). For every transcript and module detected automatically, WGCNA assigns a color based on the module membership (MM) value. An MM value is a measure of module membership which is obtained by correlating its gene expression profiles with module eigengene (i.e., the first principal component of a given module). For example, if a transcript has an MMred value close to ±1, the transcript is assigned to the red module (Langfelder & Horvath 2008). Each module is assigned a color based on the module size: ‘turquoise’ denotes the largest module, blue next, followed by brown, green, yellow and so on. The color ‘grey’ is reserved for unassigned transcripts (Langfelder & Horvath 2008). Similarly, the module ‘gold’ consists of 1000 randomly selected transcripts that represent a sample of the whole network and statistical measures have no meaning for this module (Langfelder & Horvath 2008).

After obtaining the module definitions from each comparison, we assessed how well our modules in the reference network are preserved in the population specific networks using the ‘modulePreservation’ function, which outputs a single Z-score summary. The higher the Z-score, the more preserved a module is between the reference and population-specific network. A module was deemed to be preserved if the Z-score value was above 10, an arbitrary value deemed suitable by Langfelder *et al.* (2011).

## RESULTS

### Sequencing results and mapping statistics

The dataset used for this study has been described in a previous publication by Herrmann *et al.* (2017b). Between 14 and 30 million reads were obtained for each of the 72 libraries. On average, 95.9% of the data were retained after quality control, and a mean 88.8% were mapped to the reference transcriptome. No mapping bias was observed i.e., very similar results were obtained for all genotypes. All quality and mapping metrics are available on Dryad (Herrmann *et al.* 2017a) and the raw data and experimental setup have been submitted to the ArrayExpress platform (https://www.ebi.ac.uk/arrayexpress/experiments/E-MTAB-6144). Raw reads are also available on the European Nucleotide Archive (Study ERP105101; https://www.ebi.ac.uk/ena/data/view/PRJEB23352).

### Differential expression

The intraspecific variation in transcript expression in the four populations was visualized from a read counts matrix of the 32903 transcripts using PCA (Figure 1a). A large proportion of the observed variance (19%) is explained by the first principal component (PC1). PC2 and PC3 explained 12% and 10% of the total variance, respectively. Clear population clustering is evident along PC2 for Pop.M and in Pop.J except for two genotypes (J2.1 and J2.4). However, genotypes from Pop.G and Pop.LC belong to overlapping clusters (Figure 1a and Supplementary Figure 1). No evident clustering according to experimental parameters (i.e. culture conditions, harvesting, RNA extraction batches) were visible on the PCA.

After conducting pairwise contrast analyses with DESeq2, we identified transcripts exclusively upregulated for each population when compared to all others (*p_adj_* ≤ 0.05; thereafter DETs). In total, 10,820 of 32,903 transcripts (~33%) showed significant expression differences in pairwise comparisons between populations. Of all ~33000 transcripts, 9.6%, 8.1%, 7.2% and 7.8% were population specific DETs for the populations Pop.G, Pop.J, Pop.LC and Pop.M, respectively (Figure 1b).

**Figure 1:**
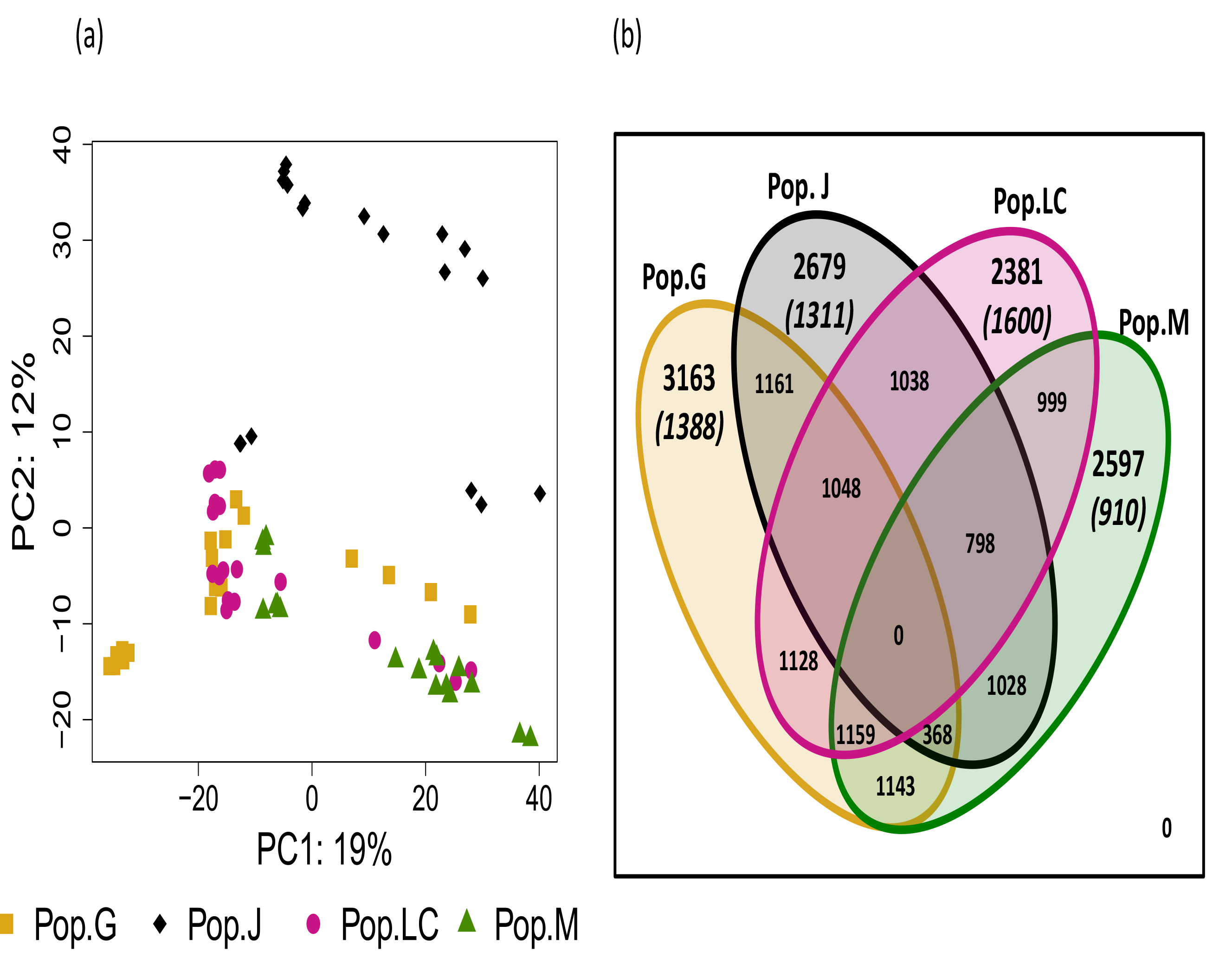
Gene expression patterns. (a) Gene expression PCA of the four sampled populations: Pop.G (Lake Greifensee), Pop.J (Jordan Reservoir), Pop.LC (Lake Constance) and Pop.M (Müggelsee). Percentages on the X-and Y-axis indicate the percentage of variance explained by each principal component. (b) Venn diagram illustrating the number of differentially expressed transcripts (DET) between the four populations. Numbers in brackets indicate the number of transcripts deviating from the neutral expectations according to the DRIFTSEL analysis.

### Role of natural selection on transcript expression levels

The DRIFTSEL multivariate approach was used to identify transcripts for which the observed differential expression could not be explained by phylogenetic distance and genetic drift alone; the alternative explanation being that the observed divergence would be attributable to selection and therefore possibly to local adaptation events. In total, 48% of 10820 differentially expressed transcripts showed greater differential expression than expected under neutrality (H-value ≥ 0.95, Figure 1b and Table S2), indicating that the observed pattern is due to local adaptation for these transcripts. Pop.LC had the highest number of DETs deviating from the neutral expectations (67% transcripts out of 2381), followed by Pop.G (~44% out of 3163), Pop.J (~49% out of 2679), and Pop.M (35% out of 2597).

### Expression variation among individuals and populations

The statistical significance of difference between group means of expression values was assessed with two ANOVAs for each transcript, one grouping the data according to genotypes and the other according to populations. For 1,897 transcripts, the means were statistically significantly different between populations but not between genotypes. The reverse was true for 7,546 transcripts. For 15,323 transcripts, the factors ‘genotype’ and ‘population’ explained the observed variation in gene expression. The remaining 8,137 transcripts had no significant *p_adj_*-values in either of the ANOVAs.

### Sequence based divergence

After applying the VariantFiltration criteria in the GATK SNP calling step, the resulting SNP set contained 414,546 variants distributed in 14,860 transcripts. These transcripts had an average of 28.2 SNPs per transcript. The vast majority (13,597 transcripts) was found to be biallelic and 1,083 transcripts were multiallelic (Table 1).

**Table 1:** Summary of SNP data. “NonDET” refers to transcripts that were not significantly upregulated in any of the pairwise contrasts.

| Population |  | all SNPs | biallelic SNPs | multiallelic SNPs |
| --- | --- | --- | --- | --- |
| Pop.G DETs | Number of transcripts | 1,369 | 1,259 | 110 |
|  | Number of SNP sites | 34,525 | 34,320 | 205 |
|  | Average number of SNPs | 25.21 | 27.25 | 1.86 |
| Pop.J DETs | Number of transcripts | 1,203 | 1,101 | 102 |
|  | Number of SNP sites | 28,252 | 28,078 | 174 |
|  | Average number of SNPs | 23.48 | 25.50 | 1.71 |
| Pop.LC DETs | Number of transcripts | 1,548 | 1,487 | 61 |
|  | Number of SNP sites | 49,451 | 49,342 | 109 |
|  | Average number of SNPs | 31.94 | 33.18 | 1.78 |
| Pop.M DETs | Number of transcripts | 1,087 | 992 | 95 |
|  | Number of SNP sites | 36,772 | 36,598 | 174 |
|  | Average number of SNPs | 33.82 | 36.89 | 1.83 |
| NonDET | Number of transcripts | 9,473 | 8,758 | 715 |
|  | Number of SNP sites | 265,546 | 264,081 | 1,465 |
|  | Average number of SNPs | 28.03 | 30.15 | 2.04 |
| Total | Number of transcripts | 14,680 | 13,597 | 1,083 |
|  | Number of SNP sites | 414,546 | 412,419 | 2,127 |
|  | Average number of SNPs | 28.23 | 30.33 | 1.96 |

A PCA was carried out based on a matrix of all biallelic SNP sites to illustrate the population structure among the four populations. Although PC1 explained the maximum variance (12%) (Figure 2a) and four distinct clusters corresponding to the populations were seen against PC2. PC2 and PC3 each explained 8% of the variance (Figure S2). PC2 clearly separated the genotypes belonging to Pop.J from the remainder of the data (Figure 2a).

**Figure 2:**
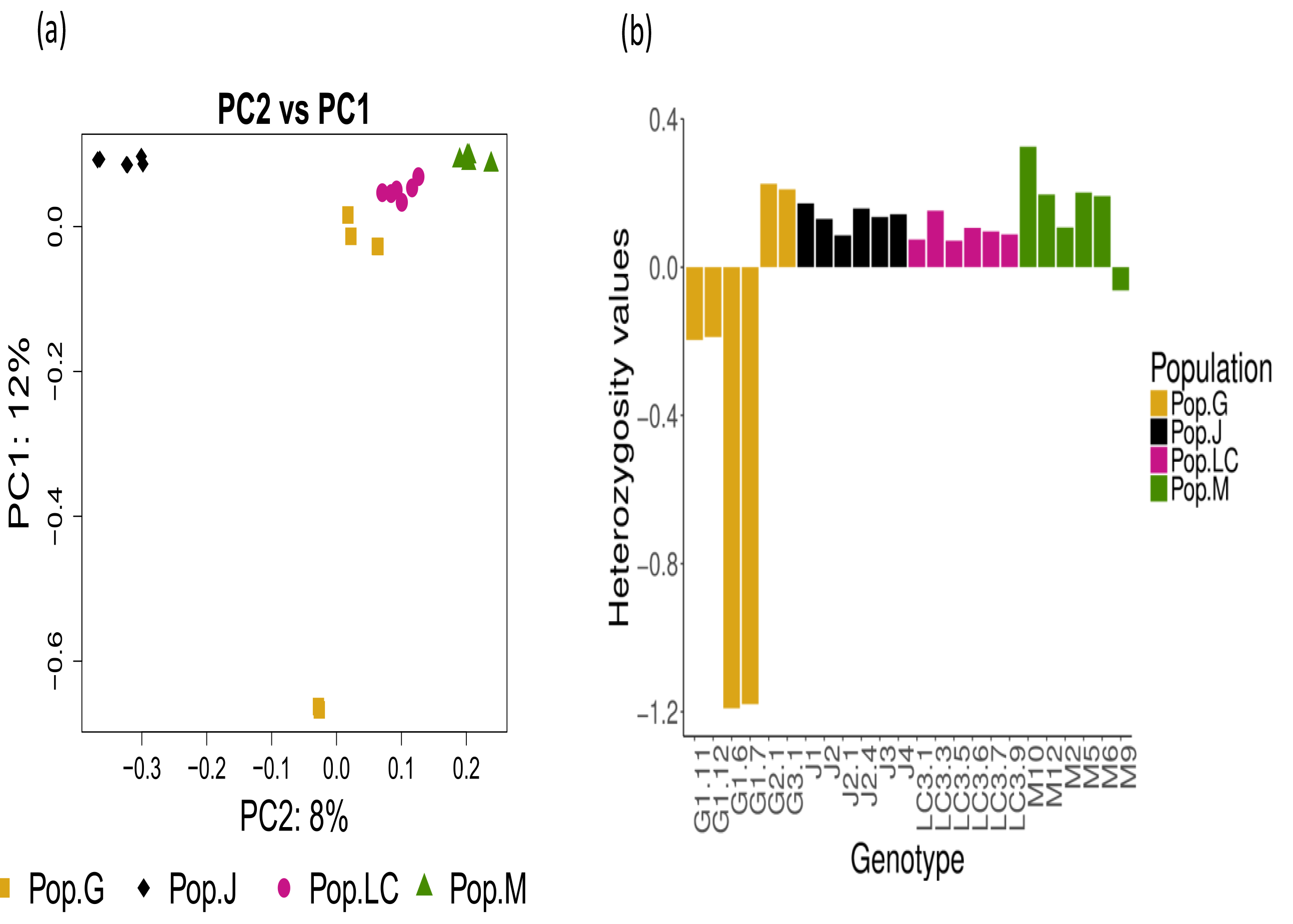
SNP patterns and heterozygosity. (A) SNP PCA of the four sampled populations: Pop.G (Lake Greifensee); Pop.J (Jordan reservoir), Pop.LC (Lake Constance) and Pop.M (Müggelsee). Percentages on the X-and Y-axis indicate the percentage of variance explained by each principal component. (B) Barplot illustrating the heterozygosity values for each genotype.

### Heterozygosity

The observed heterozygosity values ranged from −0.19 to 0.22 for genotypes from Pop.G, from 0.08 to 0.17 for those from Pop.J, from 0.07 to 0.15 within Pop.LC, and from −0.06 to 0.32 for those belonging to Pop.M (Figure 2b). Within Pop.G, four out of the six genotypes exhibited an observed heterozygosity lower than the expected heterozygosity. In Pop.J, Pop.LC, and Pop.M (except genotype M9 therein), the observed heterozygosity exceeded the expected heterozygosity; implying higher genetic variability in these genotypes.

### Sequence evolution

To assess the respective contributions of random and non-random evolutionary events on DNA sequence divergence, we calculated the Tajima’s D statistic for each transcript in the four populations. After phasing, we obtained 13,006 transcripts containing SNPs. Pop.LC had the highest number of transcripts (32.21%) with a negative D value (D < 0; *p* ≤ 0.05) followed by Pop.G (30.45% transcripts), Pop.M (29.58% transcripts) and Pop.J (29.31% transcripts). Much fewer transcripts were found to have a significant positive Tajima’s D value (Table 2): 1.26% transcripts in Pop.M, 1.20% transcripts in Pop.G, 1.13% transcripts in Pop.J and 0.66% transcripts in Pop.LC (Table S3).

**Table 2:** Tajima’s D test for selection. D < 0: number of transcripts with a negative Tajima’s D and thus likely under purifying selection; D > 0: number of transcripts with a negative Tajima’s D and thus likely under balancing selection.

| Population | D < 0 | D > 0 | Total |
| --- | --- | --- | --- |
| Pop.G | 3,961 | 157 | 4,118 |
| Pop.J | 3,813 | 147 | 3,960 |
| Pop.LC | 4,192 | 87 | 4,279 |
| Pop.M | 3,848 | 164 | 4,012 |

The LOSITAN analysis identified 782 transcripts to be under diversifying selection, 1536 transcripts under balancing selection and 113 transcripts that were under balancing and/or diversifying selection (Table S4). LOSITAN results are described in detail in (Herrmann *et al.* 2017b).

### Sequence vs. regulatory variation

The proportion of transcripts identified to be candidates for local adaptation at both sequence and regulatory level were visualized using a flow diagram (Figure 3). Among the 10,820 transcripts identified to be differentially expressed, ~46% showed signs of selection at the regulatory level according to DRIFTSEL. Of these, ~15% were identified as outliers under balancing and/or diversifying selection in LOSITAN. About 26% of these outliers had a significantly negative or positive Tajima’s D value in at least one population, which might be attributed to selection but can also stem from other evolutionary processes such as population growth, reduction or subdivision, bottleneck events and migration.

**Figure 3:**
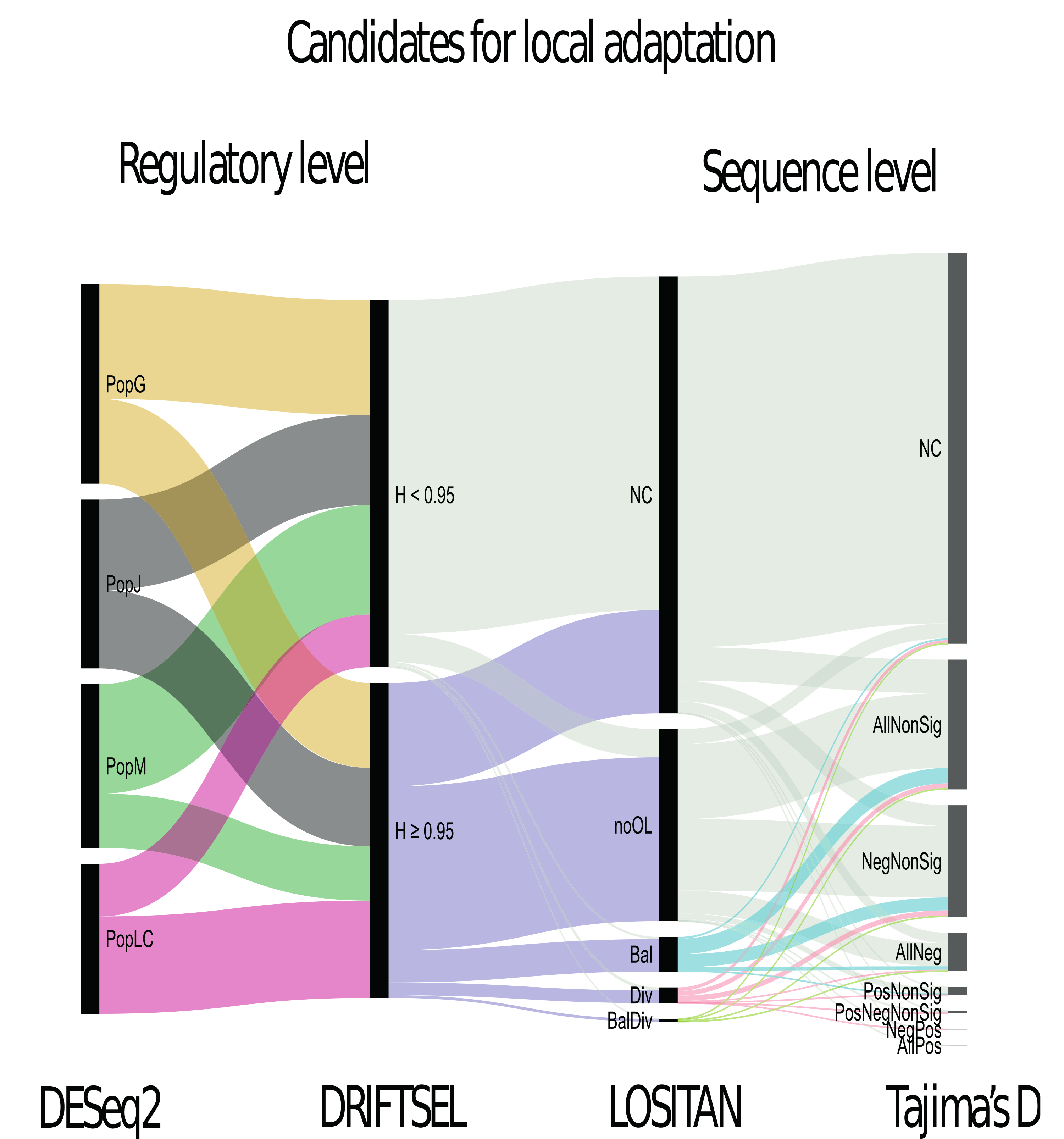
Flow diagram representing the proportion of transcripts that are candidates for local adaptation at the regulatory and sequence level. Each analysis or “step” is represented by a vertical group of black rectangle bars, called nodes. The colored areas linking the nodes are called “flows”. The **DESeq2** step contains four nodes: PopG (yellow), PopJ (black), PopLC (pink) and PopM (green), which represent the number of transcripts specifically upregulated in each of the four populations as identified by DESeq2 analysis. The **DRIFTSEL** step contains 2 nodes: ‘H.value ≤ 0.95’ (grey) and ‘H.value ≥ 0.95’ (purple). The **LOSITAN** step contains 5 nodes: ‘NC’ (grey) with transcripts without LOSITAN result (not calculated); ‘noOL’ (grey): transcripts where none of the SNPs in a transcript were identified as outliers; ‘Bal’ (cyan), transcripts containing at least one SNP that is under balancing selection; ‘Div’ (pink) transcripts containing at least one SNP under diversifying selection; and ‘BalDiv’ (pale green), transcripts containing SNPs that are under both balancing and diversifying selection. The **Tajima’s D** step contains 8 nodes. Each node classifies the transcripts according to the obtained Tajima’s D values. ‘AllNeg’ means that transcripts have a negative D value in all four populations; ‘AllPos’ means that transcripts have a positive D value in all four populations; ‘AllNonSig’ means transcripts have non-significant D values in all four populations; ‘NegNonsig’ means transcripts in the four populations have either a negative D value or a nonsignificant D value; ‘PosNonsig’ means transcripts in the four populations have either a positive D value or a nonsignificant D value; ‘PosNeg’ means transcripts in the four populations have either a positive or negative D value; ‘PosNegNonsig’ means transcripts in the four populations have either a positive or negative or an insignificant D value.

### Functional annotation

Of all transcripts, 66.5% had a BLAST hit to the nr database with an identity ≥ 50% and eval ≤ 0; 91.4% transcripts of these BLAST hits shared homology with other *Daphnia* species. Among the DETs, 70.4% met this criterion (Supplementary Figure S3a, Table S5), and 92.3% of them were homologous to *Daphnia* sequences.

We were able to predict domains for ~50% of our transcripts. Among the DETs, a slightly higher proportion of transcripts, ~53%, had known protein domains (Supplementary Figure S3b, Table S5).

For identifying *Daphnia*-specific orthologs and those that share orthology with other arthropods, the orthoMCL data was classified into six categories (as described in the Methods section). 3,058 orthology clusters (of which 1,735 clusters contained DETs) were containing exclusively *D. galeata* transcripts, 985 clusters (of which 543 clusters contained DETs) contained only *D. galeata* transcripts and *D. pulex* genes, 651 clusters (including 224 DETs) contained only *D. galeata* and *D. magna* transcripts. 3336 orthoMCL clusters (of which 1239 clusters contained *D.galeata* DETs) contained all three *Daphnia* species used in the analysis. Furthermore, 12 clusters (4 clusters containing DETs) were containing *D. galeata* transcripts along with two other arthropods (*D. melanogaster* and *N. vitripennis*). In total, 4657 clusters (1586 clusters containing DETs) contained transcripts/genes for all five species (three *Daphnia* species and two insects) used in the present study (Supplementary Figure S3c, Table S5).

### Assessment of assembly artefacts and inparalogs

In total, 3,325 DETs belonged to the 0Pop category (Figure 4a), 5,574 DETs were exclusively occurring in orthoMCL clusters without DETs from different populations (1Pop). This vast majority was thus not further analyzed with regard to paralogy and assembly artefacts. The remaining 1,921 DETs were co-occurring with DETs from other populations in 716 orthoMCL clusters. Sequence divergence was calculated for every DET pair that co-occurred in a cluster. The divergence values ranged from 0.0 to 12.0 (Figure 4b). We cannot exclude that divergence values greater than 2 between sequence pairs arose from misassemblies. However, 16,752 sequence pairs (belonging to 671 clusters) had a divergence lower than our arbitrary threshold of 2, indicating that the transcripts were highly similar in their sequence and thus might constitute inparalogs or alternative transcripts for a gene. In this case, only genomic data would allow placing the transcripts and eventually assessing their status.

**Figure 4:**
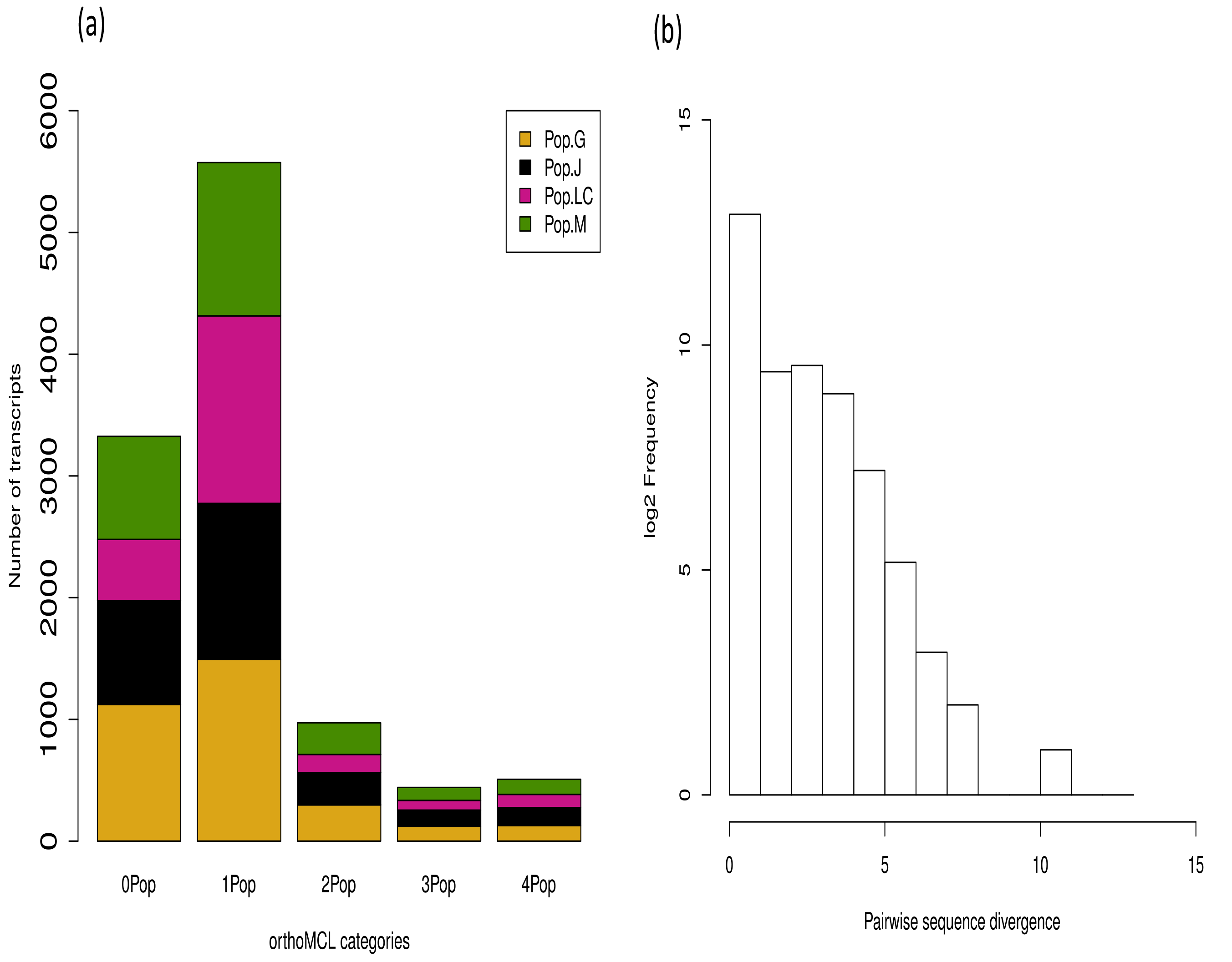
Differentiating misassembly from inparalogs. (a) Barplot showing the number of DETs co-occurring with DETs from other populations within an orthoMCL cluster. 0Pop refers to DETs not assigned to an orthoMCL cluster. 1Pop, 2Pop, 3Pop and 4Pop refer to DETs found in orthoMCL clusters containing at least one, two, three and four population(s) respectively. (b) Histogram of pairwise sequence divergence values calculated for all *D. galeata* sequences co-occurring in an orthoMCL cluster belonging to 2Pop, 3Pop and 4Pop categories.

### Gene Ontology enrichment analysis

GO enrichment analysis was performed on the candidate transcripts as identified from DRIFTSEL (Hvalue ≥ 0.95) and Tajima’s D analyses. We observed an enrichment for several metabolic processes such as ATP binding, DNA binding, microtubule binding, transporter activities and signaling pathways (Table S6) in both analyses in all population-specific sets. Specifically, in Pop.G, DRIFTSEL and Tajima’s D analysis had five GO terms in common, in Pop.J, they had one GO term in common, in Pop.LC they had four GO terms in common and in Pop.M, they had seven GO terms in common.

### Weighted Gene Co-expression network analysis

The WGCNA on 32,375 transcripts identified 29 co-expression modules (Figure 5) in the reference network (see Methods). We observed varying numbers of modules and transcripts clustered in each population-specific network (Table S7a-d). However, after assessing the conserved modules, where each population-specific network was compared to the reference network, 24 modules (out of 29) were well conserved (Zscore ≥ 10) among the populations. The conserved modules included 10,256 transcripts altogether, which is about 31% of all transcripts in *D. galeata*, with the largest module, ‘turquoise’ including 2,857 transcripts. Two modules (grey and gold) with uncharacterized and random transcripts contained 16,600 and 1000 transcripts, respectively. These results are consistent with the gene expression analysis which showed little differences between the populations.

**Figure 5:**
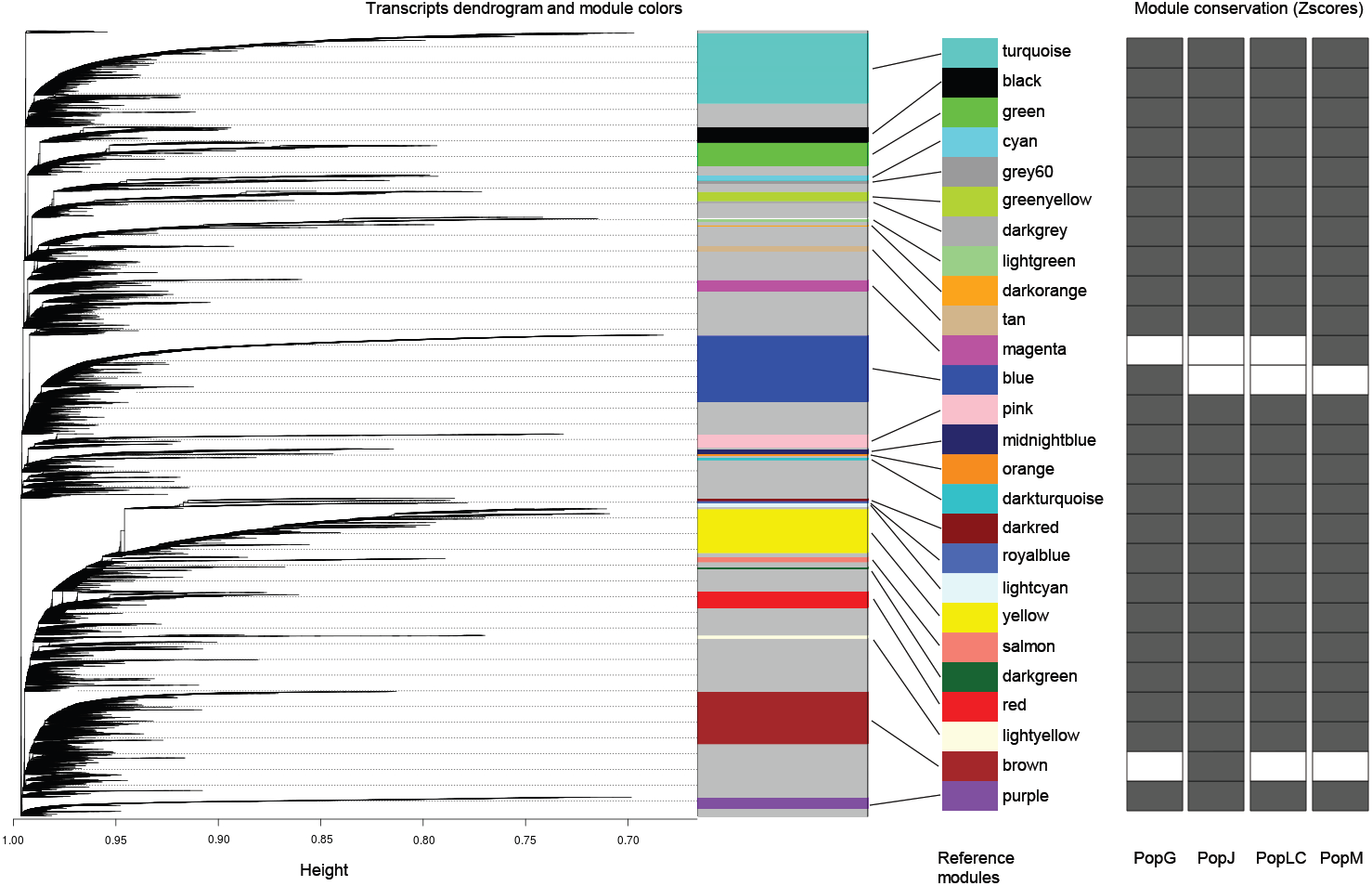
Cluster dendrogram of transcripts for the reference network in *Daphnia galeata*, with dissimilarity based on the topological overlap matrix (TOM). The co-expression modules are colored in an arbitrary way by the WGCNA package, and the size of the bar is proportional to the number of transcripts in the module. The right hand side grid represents the module conservation in each population. Modules with a Z-score ≤ 10 are shown in white and modules with a Z-score ≥10 are colored in dark grey.

## DISCUSSION

In this study, we describe an approach to distinguish between neutral and adaptive evolutionary processes at gene expression and DNA sequence level using *D. galeata* transcriptome data. We identified differentially expressed transcripts in each of the four populations. We also used the multivariate DRIFTSEL approach combining expression values and microsatellite data, to investigate the role of selection in shaping *D. galeata* differential expression profiles. Furthermore, we identified SNPs to understand the sequence level differentiation among the four populations. Finally, we annotated the functions of our candidate transcripts for local adaptation. This study is a first step towards description of polymorphisms in *D. galeata* possibly involved in phenotypic responses to environmental perturbations and as such promising candidates for future studies.

### Population divergence at the sequence level

SNPs became the absolute marker of choice for molecular genetic analysis as the mining of polymorphisms is the cheapest source for genetic variability (Taillon Miller *et al.* 1998). Our PCA analysis on SNP data revealed four clear population clusters and our results are in agreement with a highly structured population model across the transcriptome. Although two of the genotypes (G1.6 and G1.7) from Pop.G were located outside the Pop.G cluster in the PCA plot, the populations were clearly distinguished and corresponded to the four lakes sampled. This pattern might be the result of several non-exclusive phenomena: initial founder effects, isolation-by-distance and genetic drift, and natural divergent selection, since the studied populations originate from lakes located in different ecoregions.

Genetic differentiation among populations of passively dispersed aquatic invertebrates is strong in most cases, despite the dispersal probability expedited by water birds and other vectors carrying their diapausing eggs (Mills *et al.* 2007; Munoz *et al.* 2016; Ventura *et al.* 2014). Population genetic differentiation has been observed even at small spatial scales (i.e., less than 1 km) in *Daphnia* (Hamrova *et al.* 2011; Yin *et al.* 2010). Additionally, the monopolization effect, a concept based on numerous previous studies on freshwater invertebrates (De Meester *et al.* 2002; Louette *et al.* 2007; Munoz *et al.* 2008; Ortells *et al.* 2013) might reinforce the population structure resulting from initial colonization event(s). Some evidence supporting this theory has been provided by Thielsch *et al.* (2015), who showed that novel genotypes are unlikely to colonize successfully a habitat if it already harbors an established population.

All the phenomena cited above have an impact on population structure across the genome, and might mask highly diverging loci resulting from natural selection. We assessed patterns of divergence at the sequence level through neutrality tests (Tajima’s *D*). This suggested that all populations of *Daphnia* examined in this study had a substantial amount (~48% transcripts) of loci with an excess of low frequency polymorphisms (i.e., D < 0) relative to the neutral expectation. This pattern may result from positive selection, a bottleneck, or population expansion. It is consistent with previous observations in *Daphnia* from Lake Greifensee and Lake Constance (Brede *et al.* 2009) and crustacean zooplankton from Lake Constance (Straile 1998) which have all undergone historical bottleneck events. Similarly, Lake Müggelsee, a large shallow lake, has undergone severe bottlenecks due to increased turbidity and because vegetation disappeared almost completely after the 1960s (Okun *et al.* 2005). One other explanation for the excess of rare alleles is selection against genotypes carrying deleterious alleles.

Although a high frequency of rare polymorphisms was observed in our analysis, there were few transcripts (~1.7% transcripts) that had a lower frequency of rare alleles (D > 0) in the four populations; indicating that some loci are either under balancing selection (where heterozygous genotypes are favored) or under diversifying selection (where genotypes carrying the less common alleles are favored). A lower frequency of rare alleles also occurs if there is a recent population admixture (Stajich & Hahn 2005). This argument is consistent with our heterozygosity measures. Under the Hardy-Weinberg equilibrium, genotypes G2.1 and G3.1 from Pop.G, all genotypes in Pop.J and Pop.LC, and all genotypes except M9 in Pop.M show that the observed heterozygosity is greater than the expected heterozygosity, which is an indication of higher genetic variability and population admixture. Most of the genotypes in population G, as well as M9, have a much lower heterozygosity. Such low heterozygosity patterns at the individual level can be attributed to inbreeding (Keller 2002), but also due to a lack of variation in the source population, either caused by a small founder population size or a severe bottleneck during population history (Luikart *et al.* 1998). While genotype M9 from Müggelsee might be an exception, the pattern observed in Greifensee could be the consequence of inbreeding and/or low genetic variability in this population, either resulting from previous bottlenecks, or a reduced number of “founding mothers”. Further, the ecology and growth dynamics of *Daphnia* populations might exacerbate the founder effects. After an initial hatching phase from the resting eggs bank and exponential population growth in the spring, clonal selection occurs throughout the growing season (Vanoverbeke & De Meester 1997). Therefore, it is possible that only a few clonal lines contribute to the resting eggs population each year. However, while a reduced number of clonal lines might contribute to the yearly “archiving” of genetic diversity; two processes counteract the immediate diversity loss. First, the spring recruitment doesn’t only rely on eggs from the previous year but rather on a mixture (Vanoverbeke & De Meester 1997), and might even integrate overwintering clones in larger permanent lakes (but see Yin *et al.* 2014 for an overview). Second, clonal erosion doesn’t affect the same genotypes every year, leading to year-to-year heterogeneity, such as the one observed in the long term study by Griebel *et al.* (2016). Clonal erosion thus doesn’t necessarily lead to a downward spiral of genetic diversity loss, and the high stochasticity of both clonal selection and hatching ensure a preservation of the genetic diversity in every habitat.

### Gene expression variability and signals of selection

While the patterns observed at the sequence level tends to support the role of genetic drift, founder and monopolization effects in shaping the observed patterns, the results of our gene expression analysis delivered a mixed message. This was evident in the PCA based on the gene expression data, where no distinct clusters corresponding to populations are clearly visible. This observation was consistent with our network co-expression analysis which showed that the identified modules are conserved in all populations (Figure 5), with a few exceptions. The analysis of variance confirms this finding, with a relatively low number of transcripts for which the mean read counts differs significantly between populations and not between genotypes. While studies on differential expression in *Gliricidia sepium* (Chalmers et al. 1992) and *Arabidopisis halleri* (Macnair 2002) have observed substantial between population variances at the gene expression level, our results are consistent with several studies, for example, on fish (*Fundulus heteroclitus*; Whitehead & Crawford 2006a) and snails (*Melanoides tuberculata*; Facon *et al.* 2008) which showed large within-population variation. Additionally, numerous studies on life-history traits in *Daphnia* also report very high intrapopulation variability (Beckerman *et al.* 2010; Castro *et al.* 2007; Cousyn *et al.* 2001; Macháček 1991). A common garden experiment conducted on the very same clonal lines also showed a higher phenotypic variability within populations than among populations (V. Tams, personal communication). Finally, the observed relative homogeneity in the gene expression profiles might be the consequence of high selective pressure on transcription regulation or canalization (Waddington 1942). Such canalization allows for storage of cryptic genetic variation that would be uncovered in stress response assessments. However, our experimental setup was designed to avoid stress, and transcriptome characterization of the same genotypes under conditions mimicking predation, parasite or food stress, for example, might reveal a greater divergence between the populations.

Comparisons of the gene expression profiles for the four populations revealed a fair number ~8% of *D. galeata* transcripts to be significantly exclusively upregulated in one given population compared to all others. Although all populations showed similar numbers of differentially upregulated transcripts, when considering those which are probably under directional selection, the picture changed. After applying the DRIFTSEL approach, Pop.LC had the highest number of transcripts directionally selected based on their expression levels and Pop.M had the lowest number. Pop.G and Pop.J had nearly similar numbers of transcripts under directional selection. This discrepancy in the number of transcripts that are differentially expressed and those presumably under directional selection can partially be explained by parallel adaptation to contrasting environments. A study on adaptive differentiation in seagrass (Jueterbock *et al.* 2016) that compared Northern and Southern seagrass samples under thermal stress showed that natural selection was the most straightforward explanation for nearly 1% of all differentially expressed genes. For other genes that were differentially expressed in the seagrass study, parallel adaptation to different habitats was observed along both the American and European thermal clines.

### *Sequence vs. regulatory variation in* Daphnia galeata

Correlating expression profiles with sequence divergence helps to identify transcripts that are potentially under the influence of local adaptation at both gene expression and sequence level. Linking gene expression profiles with sequence polymorphisms and their associated functions aids in understanding the genetic basis of adaptation as seen in the desert adapted mouse (*Peromyscus eremicus*; MacManes & Eisen 2014) and in the Patagonian olive mouse (*Abrothrix olivacea*;Giorello *et al.* 2018). Our results revealed ~30% of the transcripts to share divergence at both sequence and regulatory level (Figure 3). There are two possible explanations for the observed differences in sequence and regulatory level variation (Hodgins *et al.* 2016). The first is that there is an increase in the rate of fixation due to transcripts under positive selection and divergence in expression patterns. For example, variation in gene expression might lead to selection for sequence variation to improve the functional role of the transcript in its altered role (Hodgins *et al.* 2016). A second explanation is that the differentially expressed transcripts may experience reduced negative selection in one or all four populations. For instance, higher transcript expression is associated with greater negative selection. Hence a reduction in transcript expression in one population compared to others may be accompanied by relaxation of selection in that population.

GO enrichment analysis on the candidates identified at the sequence (Tajima’s D) and expression (DRIFTSEL) level were enriched for metabolic and cellular processes. These findings suggest that there may be a hierarchical activation of general mechanisms of stress responses at the metabolic and cellular level. This observation is concordant to another study (Orsini *et al.* 2017) on *D. magna.* In this study, *D. magna* were subjected to several environmental perturbations and the GO enrichment analysis revealed a general stress response rather than ontologies specific to local adaptation. Since the present study is without any laboratory induced stressor, further studies in *Daphnia* subjected to one or multiple environmental stressors would be helpful in pinpointing stress specific responses. Further, no GO term annotation was available for ~31% of the transcripts, and we can therefore no reach conclusive results. This highlights the need for new and complementary resources for *Daphnia* genomics research, and a general improvement of the existing annotation.

### Gene annotation and evaluation of inparalogs

Gene annotation is quite challenging in organisms lacking reference genomes, and functional annotation then relies on the availability of transcriptomic sequences from the closest available taxon. In this study we were able to annotate 66.5% of the transcripts using BLAST analysis (Supplementary Figure S3a). However, many of the transcripts were homologous to a *D. pulex* “hypothetical protein”, likely because (i) they are similar in function to non-coding regions or pseudogenes or (ii) novel coding transcripts that are yet to be functionally characterized (Vatanparast *et al.* 2016). Furthermore, we were able to predict domains for 80% of the transcripts using Pfam analysis (Supplementary Figure S3b, Supplementary Table S4). Our orthoMCL results (Supplementary Figure S3c, Supplementary Table S4) showed that several (~45%) of the *D. galeata* transcripts were orthologous to one or all species of *Daphnia* used for comparison, indicating that the genes/transcripts have all evolved from a common *Daphnia*-specific ancestral gene via speciation. In addition to this, ~25% of *Daphnia* genes/transcripts are orthologous to two insect species (*D. melanogaster* and *N. vitripennis*). Our level of unannotated transcripts is similar to results reported from other organisms lacking extensive genomic resources, for example, from plants like field pea (*Pisum sativum;* Sudheesh *et al.* 2015), chick pea (*Cicer arietinum;* Kudapa *et al.* 2014), and winged bean (*Psophocarpus tetragonolobus*; Vatanparast *et al.* 2016). This limited our interpretation of the functional role of *Daphnia* transcripts and thereby their associations to known ecological stressors. A second issue raised when lacking a reference genome is that it might be difficult to tease apart inparalogs created by duplication events, isoforms and even misassemblies; leading to an artificially inflated number of similar sequences for each distinct gene in the transcript set. Only ~18% of the population specific DETs had one or more putative paralogs also identified as differentially expressed in at least one other population. For DETs from two or more populations that co-occurred in orthoMCL clusters, we were able to distinguish between actual paralogs (transcript pairs that had a sequence divergence value > 2, Figure 4b) and transcripts with sequence divergence value < 2.Genomic information is now required for this species in order to accurately assign transcripts to genes and correctly assess whether two different populations might indeed express different gene copies with similar functions

## FUTURE DIRECTIONS AND CONCLUSIONS

In summary, we described here an approach that combines both transcriptomic expression profiles and sequence information to understand local adaptation in *D. galeata*. Although the set of transcripts contributing to population divergence at the sequence and the expression level differ, both levels constitute alternative routes for responding to selection pressures (Pai *et al.* 2015); showing that these transcripts can contribute to local adaptation and paving way for future research. From our functional analysis, it was evident that most of our transcripts were *Daphnia* specific although they had hypothetical functions. To understand the function of the hypothetical transcripts in *D. galeata* and their response to environmental perturbations, a comparative approach using the gene expression data from numerous other *Daphnia* studies should be used. Although we noticed correlations between expression patterns and sequence divergence for the *D. galeata* transcripts, we lack genomic and phylogenetic information. This information may help “bridge the gap” for understanding the relative roles of positive or negative selection in driving coding sequence and gene expression divergence.

## Data accessibility

The raw sequence reads used for this study as well as the experimental set up for the analysis of differentially expressed genes are available on ArrayExpress (https://www.ebi.ac.uk/arrayexpress; Accession no.: E-MTAB-6144).

The raw read counts used as input for differential transcript expression, results for the pairwise contrast analysis conducted in DESEq2, and the number of variants per sample before and after filtering, number of variant sites per transcript are all available on DRYAD in Tables S10, S11 and S3-S4, respectively (https://datadryad.org//resource/doi:10.5061/dryad.p85m5). The VCF file will be made available on European Variant Archive (EVA) and accession numbers will be updated.

## Author’s contributions

SPR and MC planned the study; MC conducted the molecular work; SPR and MC designed the analysis; SPR, MC and MH analyzed the data; SPR and MC wrote the manuscript; all authors commented on results and contributed substantially to the manuscript.

## Acknowledgements

We would like to thank A. Jueterbock and O. Ovaskainen for suggestions and help on the DRIFTSEL implementation, and three anonymous reviewers for their comments on an earlier version of the manuscript. This study was financially supported by an individual grant from the Volkswagen Stiftung (to MC). This work benefits from and contributes to the *Daphnia* Genomics Consortium.

## Supporting Information

### Supporting information File 1

Table S1: Library preparation and sequencing information along with principal component coordinates for the first three axes as obtained from gene expression analysis.

Table S2: DRIFTSEL values for the differentially expressed transcripts.

Table S3: Population-wise Tajima’s D values.

Table S4: LOSITAN outlier test values to identify loci under selection.

Table S5: Functional annotation for candidate transcripts of local adaptation.

Table S6a-c: Population specific GO enrichment terms using DRIFTSEL and Tajima’s D analysis.

Table S7a-d: Number of transcripts clustered in each module as detected by WGCNA for PopG, PopJ, PopLC and PopM.

### Supporting information File 2

Figure S1 Gene expression PCA for the first three principal components Figure S2 SNP PCA for the first three principal components Figure S3a-c Pie charts showing functional annotation using BLAST, Pfam and orthoMCL analysis.

